# Identifying the functions of two biomarkers in human oligodendrocyte progenitor cell development

**DOI:** 10.1101/2020.10.27.357244

**Authors:** Haipeng Zhou, Ying He, Yinxiang Yang, Zhaoyan Wang, Qian Wang, Caiyan Hu, Xiaohua Wang, Siliang Lu, Ke Li, Zuo Luan

## Abstract

NG2 and A2B5 are important biological markers of human oligodendrocyte progenitor cells. To study their functional differences during the development of human oligodendrocyte progenitor cells to oligodendrocytes, we used cell sorting technology and obtained a large number of sterile, high-purity NG2+/- and A2B5+/- cells with high viability. Further research was then conducted via *in vitro* cell proliferation and migration assays, single-cell sequencing, mRNA sequencing, and cell transplantation into shiverer mice. The results showed that the migration ability of the cells was inversely proportional to the myelination ability. NG2 may be a marker of early oligodendrocyte progenitor cells and is conducive to cell migration and proliferation, while A2B5 may be a marker of slightly mature oligodendrocyte progenitor cells and is conducive to cell differentiation. Further, cell migration, proliferation, and myelination capacity of the negative cell population were stronger than those of the positive cell population. In summary, these results suggest that oligodendrocyte progenitor cells in the mid-stage may be more suitable for clinical cell transplantation to treat demyelinating diseases.

**Summary statement:** This research found that oligodendrocyte progenitor cells in the middle developmental stages may be more suitable for cell transplantation to treat demyelinating diseases.

## Introduction

Current clinical methods for the treatment of demyelinating disease, in addition to more mature immunotherapy, focus on cell transplantation therapy. Cell transplantation is a strategy to treat this disease by replacing the lost or damaged cell population (Alexanian et al., 2011). The transplanted cell types mainly include human oligodendrocyte progenitor cells (hOPCs) and mature human oligodendrocytes (OLs). These cells may be obtained directly from foetal and adult brain tissues (Dietrich et al., 2002; Windrem et al., 2004) or through induced embryonic stem cells (ESCs) (Izrael et al., 2007; Keirstead et al., 2005) or induced pluripotent stem cells (iPSCs) (Douvaras et al., 2014; Wang et al., 2013). Although many types of cells can be transplanted, their effects on the repair or regeneration of myelin post transplantation are variable, and these effects may be associated with the different states of the transplanted cells. Therefore, it is necessary to select a cell population that is more conducive for myelin regeneration after transplantation.

Myelin oligodendrocytes are the key cells for myelin formation. They are derived from migratory and proliferative hOPCs. Therefore, hOPCs are a potential option for cell transplantation to treat demyelinating diseases. In addition to platelet derived growth factor receptor alpha (PDGFR-α), which is a marker of hOPCs, it is well-known that chondroitin sulphate proteoglycan 4 (NG2) and ST8 alpha-N-acetyl-neuraminide alpha-2,8-sialyltransferase 1 (A2B5) are two important generally recognised markers (Hart et al., 1989; Wolswijk & Noble, 1989). Although studies have shown that both NG2-positive (NG2+) and A2B5-positive (A2B5+) cells have the ability to differentiate into oligodendrocytes *in vivo* and *in vitro* (Baracskay et al., 2007; Lyu et al., 2017), the differences in myelination effects between the two cell lineages have not been studied in depth. In addition, the sequence of expression of the NG2 and A2B5 in the development of hOPCs is also controversial. Nishiyama et al. hypothesised that the expression of NG2 occurred slightly later than that of A2B5 (Huang et al., 2020; Nishiyama et al., 2009), while Baracskay et al. suggested that NG2+ cells appeared before the expression of A2B5+ cells. Other studies have shown that as hOPCs differentiate into mature oligodendrocytes, both of these markers are downregulated (Ju et al., 2012). This dynamic change in their expression may imply that these two markers play different roles at different stages in the development of hOPCs. Therefore, elucidating the functional difference and mutual relationship between these two markers at different stages of hOPC development is the main purpose of our research.

During the preliminary research, we cultured extracted human foetal brain neural stem cells (NSCs) *in vitro* and successfully induced their differentiation into hOPCs (Lu et al., 2015). In addition, due to the fragile nature of hOPCs, MACSQuant®Tyto® (Miltenyi Biotec, Bergisch-Gladbach, Germany) was chosen for cell sorting. This new sorting method can not only yield a large number of high-purity, high-activity target cells but can also meet the gel microdroplet (GMD) levels required for clinical cell preparation, facilitating future clinical application (www.miltenyibiotec.com/local).

After obtaining the target cells, we studied the gene expression levels of these four groups of cells and the functions of the cells *in vivo* and *in vitro*. Single cell RNA sequencing (scRNA-seq) can be used to study the different subgroups of hOPCs before sorting (Feng et al., 2020; Paik et al., 2020), while RNA sequencing (RNA-seq) can be used to perform overall differential gene expression analysis and functional enrichment analysis on the sorted cell population (Liu et al., 2020; Seong et al., 2020). *In vivo* and *in vitro* cell function is assessed mainly through *in vitro* proliferation (Wang et al., 2018) and migration experiments (Crippa et al., 2019) as well as the evaluation of the myelinating ability of the sorted cells transplanted into the shiverer mouse corpus callosum (Mitome et al., 2001; Uchida et al., 2012).

This study investigated the differences in the proliferation, migration, and myelination ability of NG2+/- and A2B5+/- cells during the development of hOPCs, and revealed that the migration ability of hOPCs may be inversely proportional to their myelination ability. This provides a theoretical basis for clarifying the sequence of expression of NG2 and A2B5 during the development of hOPCs. In addition, the results of this study provide insights into the selection of cell types for cell transplantation to treat demyelinating diseases.

## Results

### hOPC identification and biomarker detection

First, we identified hOPCs using cell morphology and immunofluorescence staining. The results showed that hOPCs had the typical bipolar and bead-like appearance, and expressed PDGFR-α, NG2, and A2B5 (Fig 1A). Based on flow cytometry, the proportion of PDGFR-α+ cells was 74.13±1.74% of the whole cell population; A2B5+ cells, 25.97±0.74%; and NG2+ cells, 14.41±0.81% (Fig 1B). Further, volume uniformity of hOPCs (cell diameter <20 μm) and the expression of the three cell markers on a single cell membrane were observed (Fig 1C). scRNA-seq showed that the proportions of PDGFR-α+, NG2+, and A2B5+ cells in hOPCs were 67% (n=5309), 13% (n=1002), and 16% (n=1243), respectively (Fig 1D).

**Figure 1.**
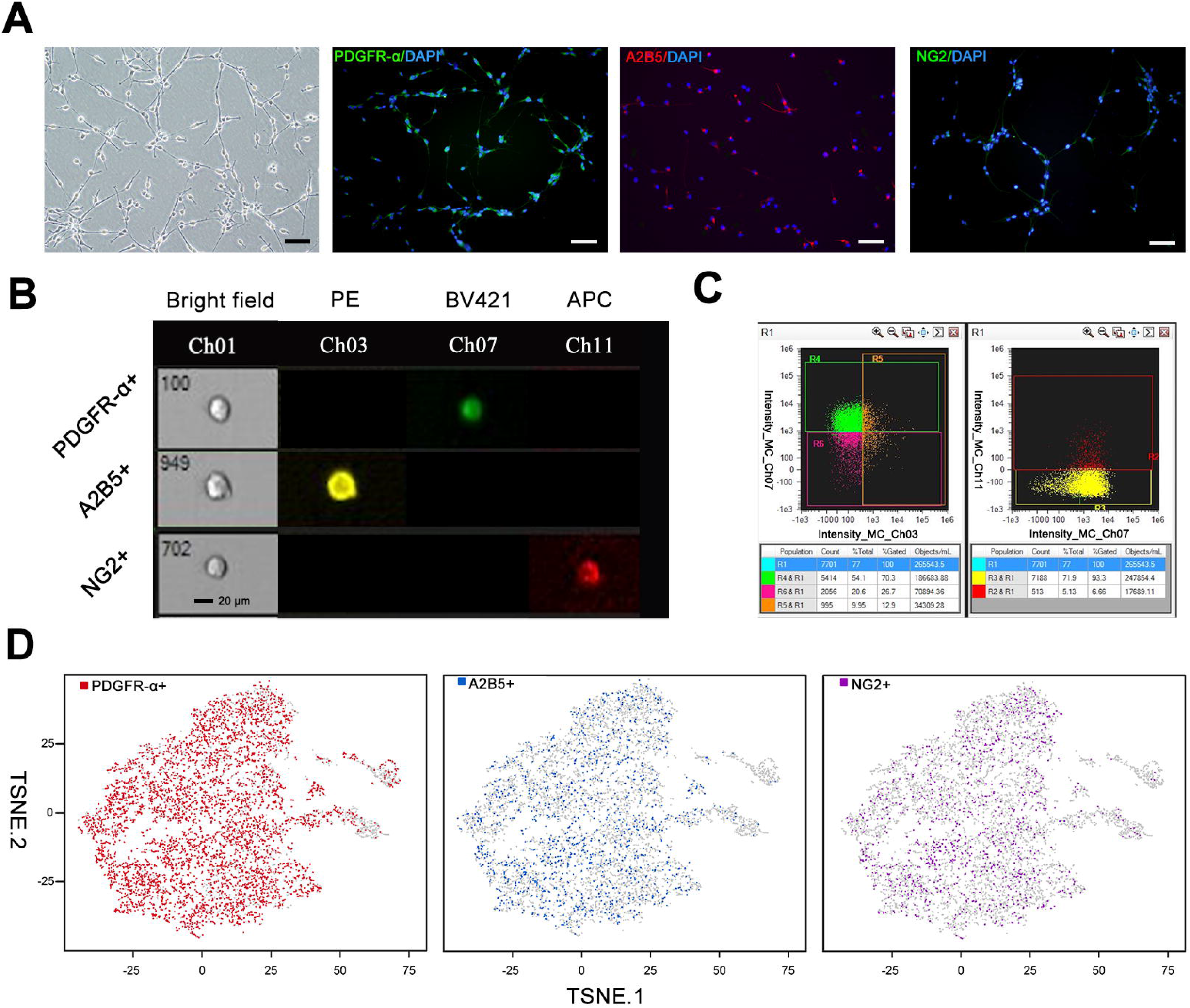
Identification of hOPC markers. A Three markers of hOPCs were identified using cell immunofluorescence staining: PDGFR-α, A2B5, and NG2. Scale bar is 200 μm. B Images of single cells detached in the presence of PDGFR-α+, A2B5+, and NG2+ cells as observed using flow cytometry. Ch01: Bright field, Ch03: A2B5-PE, Ch07: PDGFR-α-BV421, Ch11: NG2-APC. Scale bar is 20 μm. C Gating strategy to define PDGFR-α/A2B5/NG2 cells. R4: PDGFR-α+ cells, R5: A2B5+ cells, R2: NG2+ cells. D scRNA-seq of hOPCs. Maps of t-SNE of 7885 cells from high-dimensional images of hOPCs coloured according to the cell-type. hOPC, human oligodendrocyte progenitor cells; PDGFR-α, platelet derived growth factor receptor alpha; scRNA-seq, single cell RNA sequencing

### hOPC sorting and RNA-seq

To obtain NG2+/- and A2B5+/- cells, we performed cell sorting on hOPCs. After sorting, the cell viability of the sorted cells was above 98%. In the negative cell group, the proportion of A2B5+ cells dropped from 28.1 to 4.03%, and the proportion of NG2+ cells decreased from 8.22 to 1.01%. The data showed that NG2+ and A2B5+ cells were almost completely absent in the NG2- and A2B5-cell populations (Fig S1).

Further, to study the differential gene expression of the four groups, we performed RNA-seq. The results showed that, compared with A2B5+ cells, NG2+ cells had 1737 upregulated and 1225 downregulated genes. Compared with A2B5-cells, A2B5+ cells had 633 upregulated and 333 downregulated genes. Compared with A2B5+ cells, NG2-cells had 1381 upregulated and 834 downregulated genes. Compared with A2B5-cells, NG2-cells had 967 upregulated and 669 downregulated genes. Compared with NG2-cells, NG2+ cells had 466 upregulated and 352 downregulated genes. Compared with NG2+ cells, A2B5-cells had 1126 upregulated and 1400 downregulated genes (Fig 2A). Fig 2B shows a heat map of the differentially expressed genes among the four cell populations.

**Figure 2.**
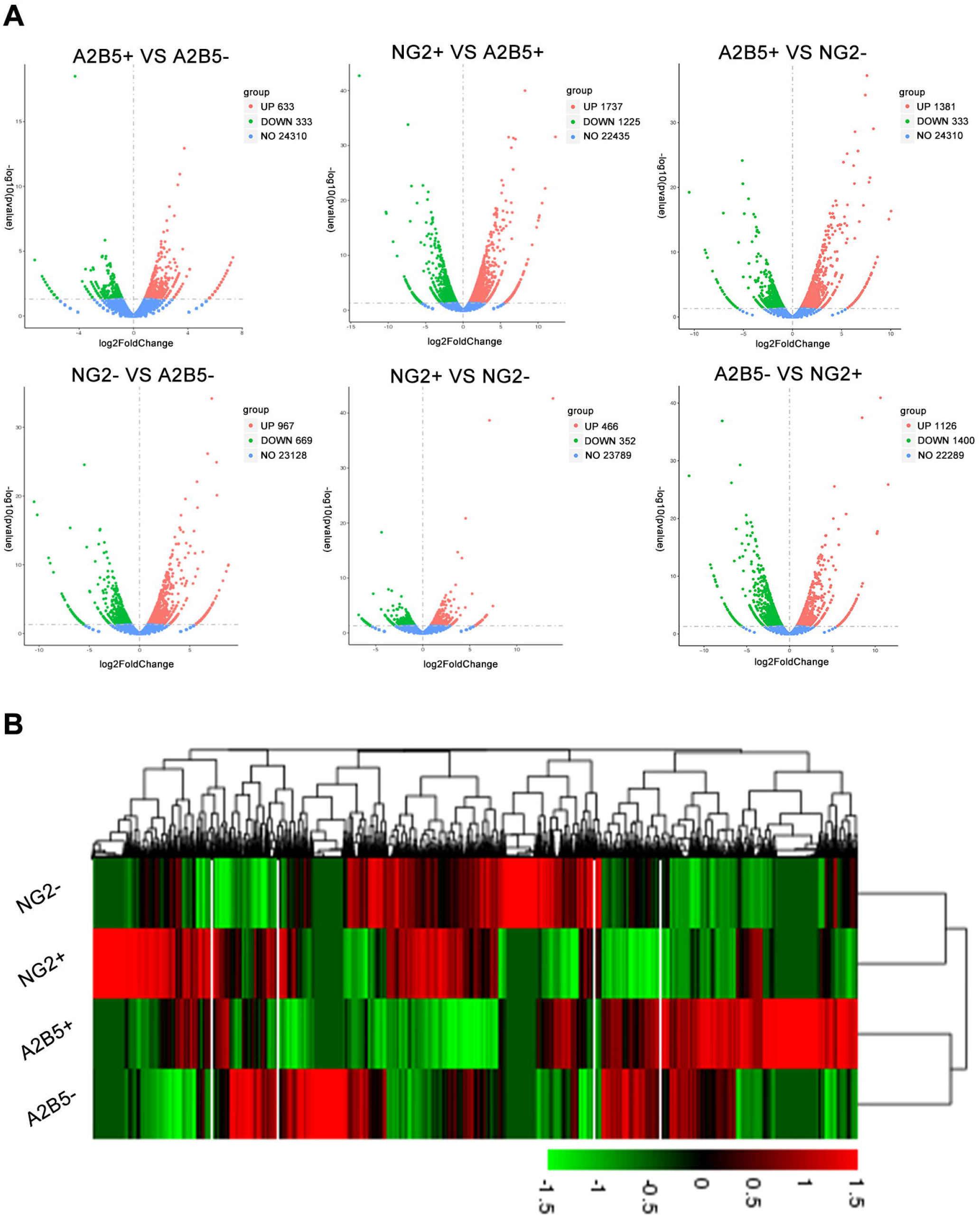
Differential expression analysis. A Volcano plot illustrating differentially regulated gene expression between the four groups of cells. Red dots indicate upregulated genes, and blue dots indicate downregulated genes. Values are presented as the log2 of tag counts. B Heat map showing the relative gene expression in NG2+/- and A2B5+/- cells. Red and green indicate upregulated and downregulated genes, respectively.

Next, we performed GO enrichment analysis on the upregulated and downregulated genes in each group. Fig 3A shows the key GO terms. BP mainly included cell proliferation (GO:0050673), migration (GO:0050673), oligodendrocyte differentiation (GO:0048709), and myelination (GO:0042552); CC mainly included extracellular matrix (GO:0031012) and axon part (GO:0033267); MF included actin-binding (GO:0003779). The bubble plots showed that in NG2+ and A2B5+ cells, the upregulated genes were mainly involved in cell migration and proliferation, while the downregulated genes were mainly involved in oligodendrocyte differentiation (Fig 3B). The results of the other groups are provided in Fig S2. In terms of cell migration and proliferation, the gene enrichment of the four groups of cells from high to low was A2B5->NG2->NG2+>A2B5+. The gene enrichment intensity for oligodendrocyte differentiation from high to low was as follows NG2->A2B5->A2B5+>NG2+.

**Figure 3.**
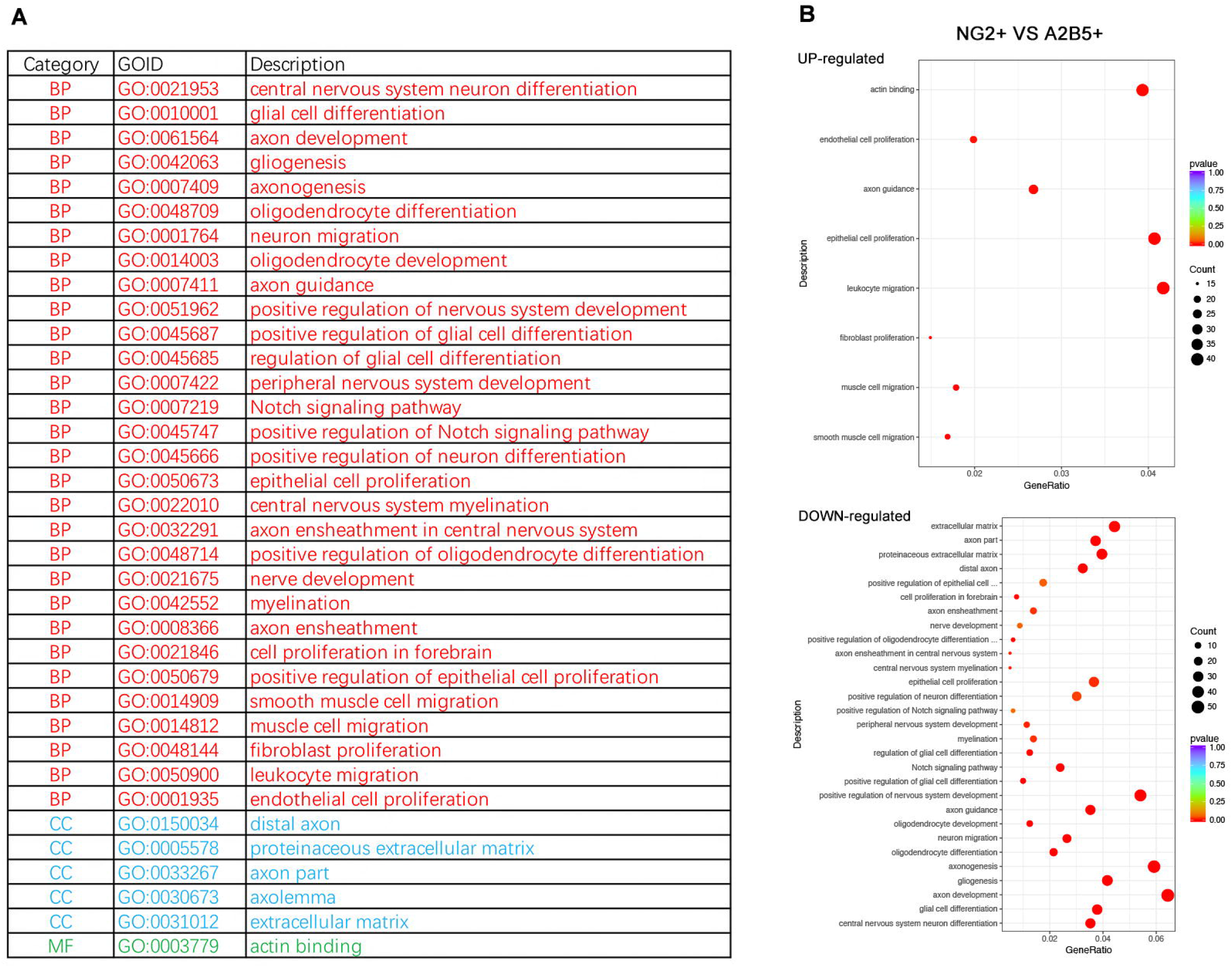
GO enrichment analysis. A Table with specific projects of BP, CC, and MF. B Bubble plots with the main enriched GO terms (NG2+ vs A2B5+) for the upregulated and downregulated genes. GO, gene ontology

### Evaluation of the migration and proliferation function of hOPCs in vitro

Next, we used the Transwell assay to evaluate the migration of the cells. After the cells had migrated for 18 h, the nuclei were stained with DAPI (Fig 4A). The results showed that the migration rates of the NG2+ and A2B5+ cell populations were 7.89±0.75% and 4.41±0.38%, respectively. The migration rates of the NG2- and A2B5-cell populations were 10.04±0.18% and 14.82±0.48%, respectively. Statistical analysis showed that the migration of NG2+ was stronger than that of A2B5+ (*, p=0.0278) and that the migration of NG2-was weaker than that of A2B5- (*, p=0.0267). The migration of A2B5+ was weaker than that of A2B5- (*, p=0.0240), and the migration of NG2+ was weaker than that of NG2- (**, p=0.0017) (Fig 4B). The proliferation assay results showed that the proliferation of NG2+ was stronger than that of A2B5+ (**, p=0.0015), while the proliferation of NG2-was weaker than that of A2B5- (**, p=0.0028). Furthermore, the proliferation of A2B5-was stronger than that of A2B5+ (**, p=0.0027), and the proliferation of NG2+ was weaker than that of NG2- (**, p=0.0079) (Fig 4C).

**Figure 4.**
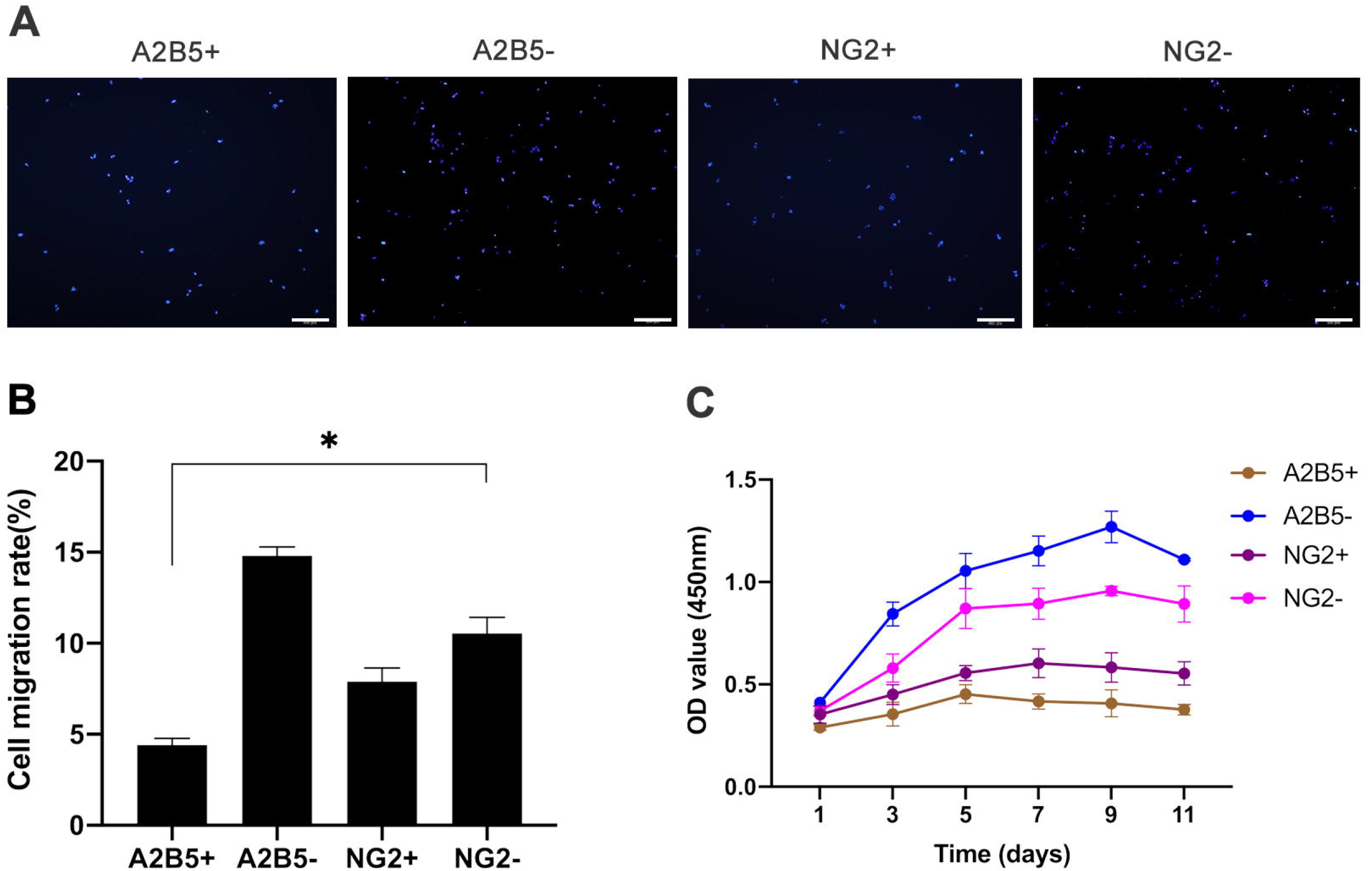
Evaluation of the *in vitro* migration and proliferation function of hOPCs. A Migration assay. Microscopic images show DAPI staining of migrating cell nuclei. Scale bar is 200 μm. B Graph showing the cell migration rate. Bars represents means. Error bars show the standard error of the mean. C CCK-8 cell proliferation assay after 1, 3, 5, 7, 9, and 11 days. hOPC, human oligodendrocyte progenitor cells, CCK, cell-counting kit.

### hOPCs were transplanted into the corpus callosum of shiverer mice

Next, we evaluated the myelinating ability of these four groups of cells *in vivo*. TEM showed that the myelin sheath of the homozygous shiverer mice did not form major dense lines or myelin compaction (Fig S3). For the shiverer mice transplanted with positive cells, although there was a small amount of myelin formation, most of the myelin sheaths were immature, the structure of the myelin was loose, and there was no compacted dense line. For shiverer mice transplanted with negative cells, the myelin sheaths were highly mature and had compacted dense lines (Fig 5A). The myelination rate analysis and G-ratio analysis of the four groups of cells showed that the NG2-cell group exhibited the best myelination effect, while NG2+ cells had the worst effect (*~****, Fig 5B, C). Finally, MBP immunofluorescence staining was performed on shiverer mouse brain tissue to further confirm myelination (Fig 5A).

**Figure 5.**
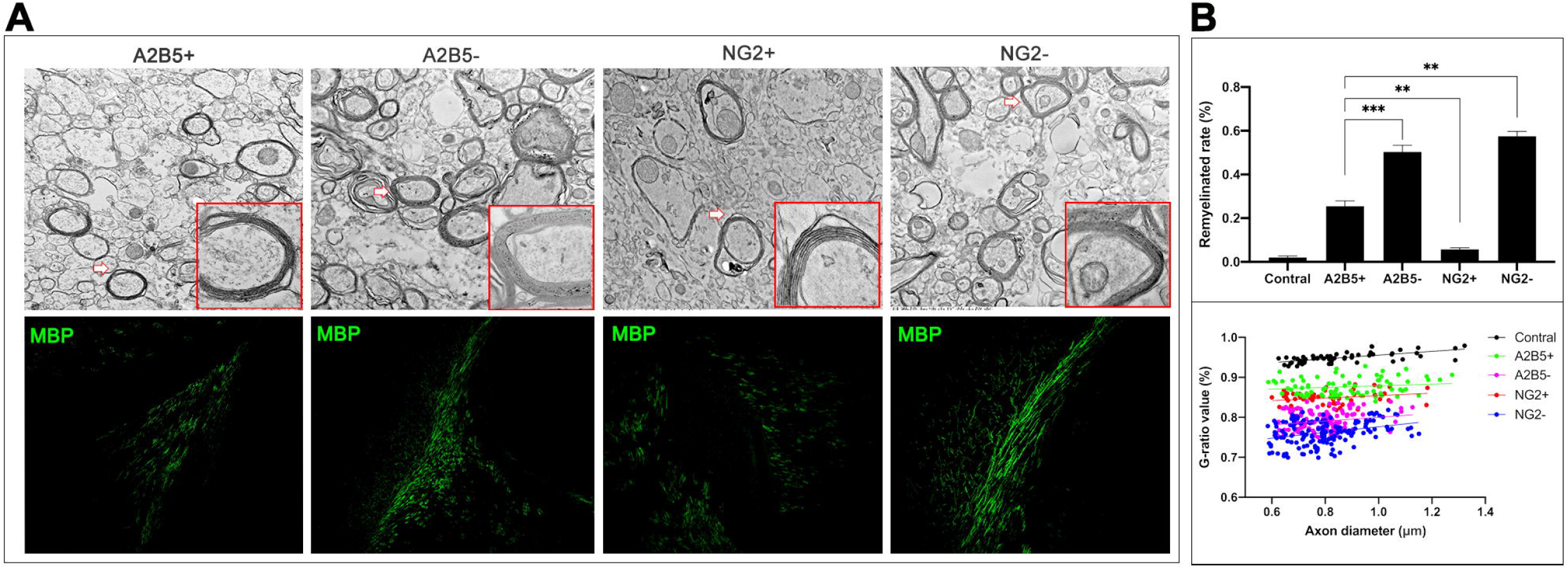
TEM and immunohistochemical analysis. A Electron micrographs showing a sagittal section through the corpus callosum of shiverer mice. Magnification: 20000X; inset 60000X. Alexa 488-labelled abundant, donor-derived MBP (green) of four transplanted cells. Scale bar is 200 μm. B Graph showing the remyelination rate of different groups and plot showing the G-ratios. Bars represent means. Error bars show the standard error of the mean. TEM, transmission electron microscopy; MBP, myelin basic protein

## Discussion

Both A2B5+ cells and NG2+ cells can differentiate into oligodendrocytes *in vivo* and *in vitro* (Diers-Fenger et al., 2001; Stallcup, 1981) and can further form a myelin structure in brain tissue, consistent with our experimental results. Other studies on the two cell populations also compared their myelinating ability. According to reports, A2B5+ cells have greater capacity to ensheath nanofibers, and NG2+ cells fail to differentiate into oligodendrocytes as quickly as A2B5+ cells (Esmonde-White et al., 2019). Our *in vivo* experiments also showed that the myelinating effect of NG2+ cells is not as notable as that of A2B5+ cells. The reason for this phenomenon may be that NG2+ cells are considered to be multifunctional cells with lineage plasticity, and NG2+ cells contain heterogeneous progenitor cells with different differentiation potentials(Schonberg et al., 2012). Because the transplanted NG2+ cells migrate extensively in the brain and differentiate into oligodendrocytes, astrocytes, and even neurons, the rate and effect of myelination of NG2+ cells are inferior to those observed for A2B5+ cells after cell transplantation.

We found that NG2-cells showed the strongest myelinating ability, while NG2+ cells showed the weakest myelinating ability. In addition, A2B5-cells showed the strongest proliferation and migration, while A2B5+ cells showed the weakest proliferation and migration. Therefore, we conclude that the migration, proliferation, and myelinating ability of the negative cell populations is stronger than that of the positive cell populations. We speculate that the reason for this may be that negative cells contain more cell types that exhibit more comprehensive and powerful functions. In addition to the above studies on proliferation-, migration-, and myelination-related functions, there is still controversy about the order of the expression of the two markers. Some studies have shown that during development, NG2+ cells originate from the subventricular region and appear before A2B5+ cells (Greenhalgh et al., 2020; Mallon et al., 2002). Other studies have shown that A2B5 is the first glial marker to appear and is expressed both on neurons and glial cells *in vivo* (Temple & Raff, 1986). More importantly, some studies have found that during the development of OLs, their myelinating ability gradually increases, while their migration ability gradually decreases (Barateiro & Fernandes, 2014). A similar observation was made in our research for NG2+ cells and A2B5+ cells, in which myelination ability was inversely proportional to migration ability. This result indicates that NG2 represents early hOPCs with strong migration ability, while A2B5 represents slightly more mature hOPCs with strong differentiation potential. Therefore, our results support the hypothesis that the expression of NG2 occurs earlier than the expression of A2B5 on cells.

In summary, we used cell sorting to sort self-made hOPCs and obtained a large number of sterile, high-purity NG2+/- and A2B5+/- cells with high viability. The feasibility of using this technology to sort relatively fragile cells is verified in this study and provides a new sorting technology reference for future clinical cell transplantation processes. Through genetic testing and cell function research, we clarified that the migration, proliferation, and myelination of a single cell population is weaker than that of a mixed cell population, and that the cell migration is inversely proportional to the myelination ability. By comparing the difference between the functions of NG2+/-, A2B5+/- cells during the development of hOPCs into OLs, we propose that hOPCs in the middle developmental stages may be more suitable for cell transplantation.

## Methods and methods

### Cultivation and identification of hOPCs

hOPCs were prepared in the Pediatric Laboratory of the Sixth Medical Centre of PLA General Hospital, China, using previously established methods of cultivation and identification. Briefly, hOPCs were induced by NSCs. The cells were cultured in a self-made medium at 37 °C in a humidified 5% CO_2_ incubator. The hOPCs were identified using immunofluorescence staining. The monoclonal mouse anti-PDGFR-α (1:250, Cat. #C2318, CST, Boston, Massachusetts, USA), mouse anti-A2B5 (1:50, Cat. # MAB1416, R&D Systems, Minneapolis, Minnesota, USA), and rabbit anti-NG2 (1:50, Cat. # ab83178, Abcam, Cambridge, Cambridgeshire, UK) antibodies were used to identify hOPCs. Cell nuclei were stained with 4’,6-diamidino-2-phenylindole (DAPI) (1:20, Cat. #28718-90-3, Sigma-Aldrich, St. Louis, Missouri, USA) for 5 min and then observed using fluorescence microscopy (IX-70, Olympus Corporation, Tokyo, Japan).

### Single-cell RNA sequencing (scRNA-seq) of hOPCs

To evaluate the gene expression of these three biomarkers, scRNA-seq was performed at the Beijing Novogene Bioinformatics Technology Co (Beijing, China). We prepared a cell suspension containing a total number of cells greater than 1×10^6^. The prepared cell suspension was quickly loaded into the chromium microfluidic chip with 3’chemistry, and a barcode of 10x Chromium Controller was attached. The cells were then subjected to RNA reverse transcription. According to the manufacturer’s instructions, a sequencing library was constructed using reagents from the Chromium Single Cell 3’v2 kit (10x Genomics, Pleasanton, California). Illumina was used for sequencing according to the manufacturer’s instructions (Illumina, San Diego, California).

### hOPC surface staining for flow cytometry and cell sorting

For PDGFR-α, A2B5, and NG2 expression analysis in hOPCs, the cells were digested and washed once with buffer (pH 7.2 PBS, 0.5% BSA) and centrifuged at 500 × *g* for 5 min at 4°C. Fragment crystallisable (Fc) receptors were blocked with normal mouse serum for 10 min at 25°C. Then, the cells were surface-stained with PDGFR-α BV421 mouse anti-human (Cat. #562799, BD Biosciences, Franklin Lake, New Jersey, USA), A2B5 PE mouse anti-human (Cat. #130-093-581, Miltenyi Biotec, Bergisch-Gladbach, Germany), and NG2 APC mouse anti-human (Cat. #FAB2585A, R&D Systems, Minnesota, USA) antibodies for 30 min at 4°C. After surface staining, hOPCs were washed once with the above-mentioned buffer and centrifuged at 500 × *g* for 5 min at 4°C. The obtained cell pellet was then resuspended in the same buffer for flow cytometry analysis and cell sorting.

### FlowSight image flow cytometric analysis

Cells were acquired using a FlowSight® imaging flow cytometer (Amnis®, part of EMD Millipore, Massachusetts, USA). Cell debris and dead cells were removed using the aspect ratio and area of the cells. Approximately 7000 cells were obtained during each analysis. Channel 7 was used to detect BV421, Channel 3 to acquire PE, and Channel 11 to detect APC. Single colour control samples were compensated using a compensation matrix (.rif) and converted to data analysis files (.daf) and compensated image files (.cif) using the same settings. The data were analysed using Ideas software, version 6.2.

### MACSQuant®Tyto® cell sorting

The fluorescent antibody-labelled cells were transferred into a MACSQuant®Tyto® Cartridge. The input sample contained 5×10^6^ hOPCs in 10 mL MACSQuant Tyto Running Buffer. Logical gating hierarchies were constructed using MACSQuant Tyto Software before sorting. First, cell debris, doublets, and dead cells were gated out and a gate was set on the target cells. The sample was sorted at 4 mL/h using approximately 140 mbar pressure. Finally, upon completion of sorting, non-sorted samples were analysed using a FlowSight® imaging flow cytometer to assess cell purity and yield.

### RNA-seq analysis

Total RNA was isolated from the four sorted populations using RNAiso Plus (Takara Bio). RNA-seq was performed at the Beijing Novogene Bioinformatics Technology Co, using the Illumina Novaseq platform. The RNA Nano 6000 Assay Kit and Bioanalyzer 2100 system (Agilent Technologies, California, USA) were used to evaluate RNA integrity. NanoPhotometer® spectrophotometer (IMPLEN, California, USA) was used to check the purity of the RNA. Clustering and sequencing (Novogene Experimental Department) were conducted according to the manufacturer’s instructions. The index-coded samples were clustered using a cBot Cluster Generation System and TruSeq PE Cluster Kit v3-cBot-HS (Illumina). Finally, the library preparations were sequenced using the Illumina Novaseq platform and 150-bp paired-end reads were generated.

### Differential expression analysis and gene ontology (GO) enrichment analysis

First, we used the edgeR program package with one scaling normalization factor to adjust each sequenced library for read counts, and then performed differential gene expression analysis. p values were adjusted using the Hochberg and Benjamini method. Differentially expressed genes were analysed for GO enrichment by the clusterProfiler R software package, and the bias in gene expression was corrected. GO terms included are as follows: biological process (BP), cellular component (CC), and molecular function (MF). When the corrected p value was <0.05, differentially expressed genes were considered to be significantly enriched in the GO terms.

### Cell migration assay

Cell migration was also measured in a Transwell filter with 8-mm pores (Corning, Tewksbury, Massachusetts, USA). Inserts were coated with fibronectin human protein (Cat. #33016015, Thermo Fisher Scientific, Waltham, Massachusetts, USA). Sorted cells (2×10^4^ cells), in 200 μL of the medium, were loaded to the upper chambers and 500 μL of the hOPC medium was added to the chamber. After incubation for 18 h at 37°C, a cotton swab was used to wipe the cells that had not migrated onto the upper surface of the chamber. Next, the migrated cells were fixed with 4% paraformaldehyde and stained with DAPI (1:20). Images were acquired using a fluorescence microscope. The migrated cells were quantified using ImageJ software by analysing cell nuclei from at least five randomly selected fields per chamber. The cell migration rate was calculated as follows: cell migration rate (V1%) = N1/N2 × 100%, where *N1* refers to the initial cell seeding number, and *N2* refers to the number of migrated cells. The experiment was repeated thrice independently in the laboratory.

### Cell proliferation assay

Cell proliferation of different cell populations was evaluated using the cell counting kit-8 (CCK-8) assay (Cat. #CK04, Dojindo, Kumamoto, Japan). hOPCs (6000 cells/well) were seeded into 96-well plates and cultured for 24 h at 37°C and 5% CO_2_. The same volume of hOPC medium was added to the blank control group, and six parallel wells were set for each group without any cells. At different experimental time points (1, 3, 5, 7, 9, and 11 days), 10 μL of CCK-8 solution was added to the wells and incubated at 37°C for 2 h. Six wells were set for each experiment at each time point. The absorbance at 450 nm was measured with a microplate reader (BioTek, Winooski, Vermont, USA). At least three experiments were performed, and each was tested in triplicate. The experiment was repeated thrice independently in the laboratory (Fu et al., 2017).

### Cell transplantation in shiverer mice

Homozygous shiverer mice (The Jackson Laboratory, Bar Harbor, Maine, USA) were maintained at the Sixth Medical Centre of PLA General Hospital in a specific pathogen-free environment. All animal experiments were performed according to protocols approved by the Sixth Medical Centre of PLA General Hospital Animal Care and Use Committee (Application No. SCXK-2012-0001). New-born pups were transplanted within a day after birth. Different cell populations (1×10^5^/1.5 μL) were injected bilaterally into the corpus callosum using a mouse brain stereotaxic apparatus (Stoelting, Chicago, Illinois, USA). At the same time, untransplanted homozygous shiverer mice (n=6) were randomly selected as the control group. The postoperative pups were returned to their mother and were weighed every day. After weaning, each mouse was reared separately. At three months of age, the mice were anaesthetised with 1% pentobarbital sodium. Then, mouse brain samples were perfused with PBS, and fixed with 4% paraformaldehyde (Cat. #MFCD00133991, Thermo Fisher Scientific, Waltham, USA).

### Transmission electron microscopy (TEM)

After the corpus callosum was separated, it was immediately fixed in 2.5% glutaraldehyde at 4°C, and then fixed with 1% osmic acid for 2 h. After embedding with epoxy resin, tissue samples were sliced into ultrathin sections using a PowerTome X Ultramicrotome (RMC products by Boeckeler, Tucson, Arizona, USA). The samples were then stained with uranyl acetate and lead citrate, following which, the structure of myelin was observed using TEM (H7650-B, HITACHI, Tokyo, Japan). Then, a field of view was randomly selected for each mouse, and the number of myelin sheaths in the photo was quantified. Finally, the remyelination rate and G-ratio were calculated.

### Tissue immunofluorescence staining

The mouse brain samples were fixed in 4% formaldehyde overnight and immersed in 25% sucrose solution for 3 days. Serial frozen sections (5μm thick) were obtained after embedding at optimal cutting temperature. Myelination was identified using immunohistochemical staining. Cryosections were rewarmed at 37°C for 1 h and then rehydrated in 0.03% Triton X-100 in PBS for 45 min. Blocking was performed in 3% normal goat serum (Cat. #G9023, Sigma-Aldrich, St. Louis, USA) for 45 min. The rat anti-myelin basic protein (MBP) (Cat. #Ab7349, Abcam, Cambridge, UK) antibody was diluted in blocking solution at 1:200 and incubated overnight at 4°C. After washing with PBS the next day, the brain tissue was incubated with the secondary antibody (Alexa Fluor® 488 donkey anti-rat IgG, 1:500, Cat. #Ab150153, Abcam, Cambridge, UK) in a dark room for 1.5 h, and then washed with PBS. A fluorescence microscope (IX-70, Olympus Corporation) was used to observe the fluorescently labelled samples, and the mean optical density of MBP was analysed using Image Pro Plus (version 6.0).

### Statistical analysis

The experimental data in this study were analysed using GraphPad Prism version 8 and SPSS version 22.0. Differences were considered statistically significant at p<0.05. Statistical significance was determined using independent samples Student’s t-test or one-way ANOVA followed by Bonferroni tests. All experimental data are expressed as mean±SD.

## Acknowledgements

The authors appreciate the technical support and other help from Leping Zhang, Jing Zang, Chang Liu, and Qian Guan. We would like to thank Editage (www.editage.cn) for English language editing.

## Author contribution

HPZ, YXY, and ZL designed the experiments. HPZ, YH, YXY, ZYW, QW, CYH, XHW, SLL, KL, and ZL performed the experiments. HPZ, YH, and YXY analyzed the data. ZYW, QW, CYH, XHW, SLL, KL, and ZL contributed materials/reagents/analysis tools. HPZ wrote the manuscript.

## Data availability

All data generated or analyzed during this study are included in this published article. All of the raw data for this study will be forthcoming.

